# Fossil-based analyses of clades’ diversification patterns require taxonomic expertise and appropriate methodology

**DOI:** 10.64898/2026.02.27.708174

**Authors:** Guillaume Guinot, Sylvain Adnet, Gilles Cuny, Iris Feichtinger, Kenshu Shimada, Mikael Siversson, Charlie J. Underwood, Romain Vullo, David J. Ward, Fabien L. Condamine

## Abstract

Estimating deep-time diversification patterns and the establishment of extant biodiversity represent major challenges in macroevolution. Fossil record data provide essential information to address these topics, but their heterogeneous temporal and geographical distributions require using analytical approaches to process these data. Gardiner et al.^1^ (hereafter GEA) used a deep-learning model^2^ and a fossil-occurrences dataset^3^ to estimate neoselachian richness over the last 145 myr.

**Results and Discussion:** GEA^1^ found that neoselachian diversity increased throughout the Cretaceous, was little impacted by the Cretaceous-Paleogene (K/Pg) mass extinction (~10% species loss), and peaked in the mid-Eocene but declined until the Present. While the Cretaceous increase in neoselachian richness is well known^4^, the other findings of GEA^1^ are at odds with current knowledge. With the exception of lamniform sharks, the perceived decrease in species richness in the recent past is most likely due to a drop in available fossil record data combined with difficulties in identifying extant species in the fossil record^5^. Similarly, all previous analyses of the impact of the K/Pg mass extinction on elasmobranch diversification have reported high extinction rates, a marked diversity drop, and delayed recovery^6–7^, despite heterogeneity across clades, ecology, and geographical distribution^7^. Taking the K/Pg as an example, we demonstrate that the discrepancies between GEA^1^’s results and current consensus is most likely due to a combination of incomplete, unverified, and incorrect fossil-occurrence data with inappropriate methodology.

## Fossil-occurrences dataset and taxonomic expertise

GEA’s^1^ fossil-occurrence dataset is currently available as a preprint^3^ and was used in a previous analysis^8^. This dataset combines data extracted from the Paleobiology Database (http://www.paleobiodb.org/) and Shark-References (https://shark-references.com/), an online compendium of chondrichthyan-related literature. This dataset is presented as extensively curated, with taxonomic nomenclature and occurrences’ age used deemed to be valid^1^. After filtering, the dataset contains 24,464 occurrences at the genus level and 18,328 at the species level.

GEA^1^ found an absence of pronounced K/Pg extinction, contradicting current consensus based on various methods and data^6–7,9^. We consequently reexamined their dataset, focusing on the survivors and victims of the K/Pg event. From their main dataset, we selected species and genera with occurrences distributed across the K/Pg boundary (survivors) and those having the younger age of their last occurrence at 66 Ma (victims). We then checked the taxonomy, verified the occurrences of survivor taxa, and plotted their distribution through time to identify outliers and discrepancies. We identified 146 victims and 81 survivors for species (genera: 51 victims, 100 survivors). Our verification revealed that 83.95% (68/81) of species identified as survivors in GEA’s dataset are in reality restricted to the Paleogene or Maastrichtian (54.84% for genera: 51/93 plus seven duplicates) (Supplemental Material). These incorrect occurrences strongly impact the conclusions drawn by GEA^1^ and are mainly affected by erroneous taxonomic attributions and incorrect age assignments.

### Taxonomic issues

One of the most pervasive and problematic taxonomic flaws is the treatment of uncertain nomenclature (i.e., taxon names containing “cf.”, “aff.”, “?”), which indicates a species that shares resemblance or affinity with, but is not identical to, the named species. In GEA^1^’s dataset, all such occurrences were assigned to the named species they were originally tentatively referred to. For example, the Danian occurrence originally reported as *Squatina* aff. *cranei* is unjustifiably treated as *Squatina cranei*, although all other occurrences of this species (including the type specimens) are ~30 myr older. Such taxonomic decisions constitute a significant number of outliers in occurrence distribution through time and contribute to the artificially high number of K/Pg survivors. We identified 1,821 occurrences (7.4%) originally reported in open nomenclature but assigned to a named species by GEA^1^. Such occurrences should instead be treated on a case-by-case basis or be ranked at higher taxonomic level.

Another major taxonomic issue in GEA^1^ is the inclusion of a tremendous number of occurrences considered valid although not based on any evidence (i.e., no illustration, specimen number, nor description). While such occurrences represent an evident source of error and should be discarded as their identification is unverifiable and their taxonomy impossible to update, they represent 39% (9,561) of occurrences in GEA^1^’s dataset. These two major errors produced Paleogene occurrences for many decisively Cretaceous taxa (e.g., *Onchosaurus, Squalicorax kaupi*) and Cretaceous occurrences of definitive Cenozoic taxa (e.g., *Isurus, Myliobatis dixoni*) (Figures S1-4).

Although taxonomic verification was made difficult by the absence of author names for taxa, we identified seven survivor genera representing duplicates (e.g., *Cretolamna*/*Cretalamna*) or disused names (e.g., *Oxyrhina*). Outdated taxonomy, with K/Pg survivor species included in a Cretaceous genus, or taxonomic misunderstanding (e.g., Cretaceous “*Dasyatis* Rafinesque” assigned to the Ypresian species *Dasyatis rafinesquei*), further artificially increased the number of survivors. We also identified serious errors in the attribution of genera to higher taxonomic levels (e.g., *Potamotrygon* species within Rajiformes instead of Myliobatiformes), affecting order-level analyses. The use of the bibliographical database Shark-References for taxonomy cannot replace careful examination of the data.

### Age issues

Another large part of outliers in the stratigraphic distribution of species crossing the K/Pg boundary relate to errors in the age assignment of some fossil collections. For example, the age of locality #1155 (Sillon X+Dan at Sidi Daoui, Morocco) is considered Maastrichtian-Danian in the dataset, whereas it is well known that this horizon corresponds to the mixing by bioturbation of Maastrichtian and Danian layers. Assigning a Maastrichtian-Danian age to the corresponding 22 species in the dataset led to the creation of outliers for typical Cretaceous and Danian species, which artificially become survivors of the K/Pg. This underscores the importance of carefully considering geological settings when compiling fossil datasets. Finally, we identified 14 essential species missing among K/Pg victims and omission of key occurrences, including those representing the type locality of obligate Maastrichtian or Paleogene species. This results from the flawed choice of excluding literature older than 1970 from references extracted from Shark-References.

### Choice of appropriate methodology

GEA^1^ aimed to assess neoselachian diversity through time and to identify what is referred to as the “*magnitude of species loss*” or “*change in species richness*” between identified “*phases*”. They applied a deep-learning model^2^ informed by the distribution of fossil occurrences to estimate paleodiversity through time (accompanied by SQS, PyRate, *Divvy*) and predict (unobserved) species richness. This approach overlooks underlying speciation and extinction rates, except for the K/Pg event (PyRate). We argue that analyzing the evolution of species richness through time cannot be done without considering associated speciation and extinction rates, regardless of the method used. A multitude of evolutionary histories can produce similar diversity trajectories, and isolated richness predictions across diversification phases are uninformative unless rates are provided.

Focusing on the K/Pg event, GEA^1^ found a 9.7% ±9.1% (32% ±2% based on raw data) loss in neoselachian species richness from the Maastrichtian to the Danian. This contrasts with estimates of species extinction (62.63%) found through Bayesian modelling of fossil-occurrence data^6^. GEA^1^ explained this discordance based on their “increased” dataset and their analytical approach. Among the major issues GEA^1^’s dataset suffers from, the 39% of unverified occurrences produces a higher number of occurrences, but it has been shown that an amplified number of occurrences does not impact estimated diversification rates at constant number of species^10^. Thus, differences in dataset content cannot fully explain the divergence in extinction magnitude.

We contend that the methodology used to calculate species loss explains these differences. Previous estimates of the extinction magnitude of the K/Pg^7^ were made by computing the number of species going extinct (*e*) over standing diversity (D), thus providing conventional extinction percentages (*e*×100/D)^9^. This was further conservatively calculated over the 800-kyrs extinction interval, and not across the whole Maastrichtian. GEA^1^ opted for calculation of net change (%) in species richness between the Maastrichtian and Danian (and repeated this for the six identified phases). This simply compares absolute richness between two time bins regardless of the number of apparitions/extinctions creating the observed difference. This naive and unusual approach is meaningless in paleobiodiversity analyses as it fails to illuminate extinction magnitude. Lack of data on victim/survivor lineages estimated in GEA^1^ prevents conventional comparative calculation of the extinction magnitude found in their analyses. However, using the raw and flawed fossil-occurrence data provided in GEA^1^, we find an extinction magnitude of 60.55-64.32% for species (genera: 30.83-33.77%), while GEA found a change in taxonomic richness of −32% ±2% for species (genera: −23% ±1%). Once dubious occurrences and taxonomy were corrected, the magnitude of the K/Pg extinction rises to 90.29-93.53% for species (genera: 63.73-65.29%) using the conventional method. This demonstrates that GEA’s methodology (richness without rates, flawed estimate of ‘species loss’ magnitude) is inappropriate to tackle the questions addressed.

## Conclusion

The evolutionary history of neoselachians is long and complex, gleaned from their rich fossil record with over two centuries of taxonomic history. Given their importance in marine ecosystems through geologic time, it is critical to understand their diversification history, but this requires appropriate methodology and fossil-occurrences data curated with taxonomic expertise. Failing to do so produces spurious results and conclusions. By underestimating the magnitude of the K/Pg mass extinction, GEA^1^ convey the misleading impression that this clade—currently under severe threat—is resilient to extinction, although evidence indicates that it experienced substantial losses in the past^4,6,7^. Considering the numerous and serious flaws GEA^1^’s dataset contains, we discourage its use in fossil-based diversification analyses.

## Supplemental Information

Supplemental information including procedures, figures, tables, and references can be found online.

