## Supplementary Information file for "Fossil-based analyses of clades’ diversification patterns require taxonomic expertise and appropriate methodology"

#### Supplemental procedure

##### *Verification of the dataset*

Gardiner et al. (2026) found that neoselachian diversity increased throughout the Cretaceous and was little impacted by the Cretaceous-Paleogene (K/Pg) extinction event (~10% species loss). They further found that the maximum neoselachian diversity peaked in the mid-Eocene and declined until the Present. While the Cretaceous increase in neoselachian richness is known from previous global and clade-centered analyses (Guinot et al. 2012; Guinot & Cavin 2016; Brée et al. 2022; Condamine et al. 2019; Marion et al. 2025, 2026), the other findings of Gardiner et al. (2026) are at odds with current knowledge. One of them is the absence of marked extinction at the K/Pg boundary. Because this statement in particular contradicts current knowledge based on various methods and data that provided solid evidence for both elasmobranchs (Kriwet & Benton 2004; Guinot & Cavin 2016; Guinot & Condamine 2023) and metazoan in general (Sepkoski 1984; Stanley 2016), we chose to explore the dataset of Gardiner et al. (2026) by mainly focusing on the survivors and victims of the K/Pg event, although some issues led us to proceed to verifications across the entire dataset.

All 24,464 occurrences used in Gardiner et al.'s analyses were provided in their Data S1 and correspond to 'filtered data' from the unpublished FINS dataset (Kocáková et al. 2025). From the dataset in Data S1, we first selected the species with occurrences distributed at each side of the K/Pg boundary (66 Ma) (i.e., the survivors) and the species having the younger age (column *min_ma*) of their last occurrence at 66 Ma (i.e., the victims). We then checked the taxonomy and the occurrences of the survivor species, and plotted their distribution through time to identify potential outliers around the K/Pg boundary and illustrate discrepancies. We repeated this procedure for genus-level data (column "*rank*" including both *species* and *genus* entries). We restricted our occurrence verification to the youngest occurrences for typical Cretaceous taxa indicated as surviving the K/Pg extinction and to the oldest occurrences of typical Paleogene taxa, which are considered present in the Cretaceous.

We identified 146 victims (Table S1) and 81 survivors (Table S2) for species and 51 victims (Table S3), and 100 survivors for genera (Table S4). Among the 81 species identified as survivors in Gardiner et al.'s dataset, we found that 83.95% (68/81) are in reality either restricted to the Paleogene or to the Maastrichtian (Table S2). Among the 100 genera identified as survivors in Gardiner et al.'s dataset, seven are duplicates of other genera (*Cretolamna/Cretalamna, Notidanus/Hexanchus, Rhinobatos/Rhinobatus*) or correspond to disused names (*Eorhincodon, Hypolophus, Oxyrhina, Orthacodus*) and 54.84% of the remaining (51/93) are actually either restricted to the Paleogene or to the Maastrichtian (Table S4).

We extracted the occurrences of each survivor species of dubious temporal distribution by subsetting Gardiner et al.'s dataset using the species name in the "*accepted name*" column. We repeated this procedure for genera, using the column "*genus*" for filtering by genus name. We then plotted the occurrences per taxon using the geom_violin function of the *ggplot2* package (Wickham 2016) in R (R Development Core Team 2025). We plotted the younger age ("*min_ma*" column) of each occurrence for typical Cretaceous taxa and the older age ("*max_ma*" column) of each occurrence for typical Danian taxa. Corresponding plots are available in Figures S1-4.

For each survivor taxon with dubious temporal distribution, we identified their minimum age (youngest age of occurrences in the "*min_ma*" column) and maximum age (oldest age of occurrences in the "*max_ma*" column), the number of outlier occurrences making the taxon cross the K/Pg boundary, their occurrence number as entered in Gardiner et al.'s Data S1, corresponding references, the "*evidence*" status, and potential taxonomic and/or age issues we identified (Tables S2, S4).

Considering the large number of outliers represented by occurrences where taxon names were originally reported in open nomenclature but treated as nominal species in the dataset, we searched for the proportion of such occurrences across the entire Dataset S1. We extracted the occurrences corresponding to taxa left in uncertain/open nomenclature (i.e., names containing "cf.", "aff.", "?"), by searching for "cf.", "aff.", and "?" in the column '*identified_name*' of Gardiner et al.'s Data S1. We identified 1,821 occurrences (7.4%) originally reported in open nomenclature (296 as "?", 1,059 as "cf.", and 466 as "aff.") but assigned to a named species in the dataset (column "*accepted_name*").

As it appeared that a large number of outlier occurrences of survivor taxa were based on reports made without evidence (i.e., no illustration, no specimen number, no description in source publication), we searched for the amount of such occurrences across the entire Gardiner et al.'s dataset. We filtered the column "*evidence_validation*" with the entry *no_evidence* and identified 9,561 unverified occurrences among the 24,464 occurrences contained in the dataset (39.08%). Such unverified occurrences can originate from literature such as conference abstracts (Dutheil 1996) or species lists provided in geochemistry analyses (van Baal et al. 2013).

We found further taxonomic issues such as where K/Pg survivor species are included in a Cretaceous genus (e.g., *Palaeotriakis curtirostris* included in the Cretaceous *Paratriakis*), or misunderstanding of the taxonomy (e.g., Cretaceous occurrence reported as "*Dasyatis* Rafinesque" assigned to the Ypresian species *Dasyatis rafinesquei*), further artificially increased the number of survivors (Tables S2, S4).

We did not check in-depth erroneous attributions of genera to orders but spotted many instances where such incorrect attributions were present in the dataset and not supported by any published hypothesis: seven species of *Potamotrygon* (Myliobatiformes) within Rajiformes; four species of *Squatirhina* (Rhinopristiformes) within Rajiformes; two species of *Belemnobatis* (Rhinopristiformes) within Rajiformes.

We checked for age assignment of some fossil collections around the K/Pg boundary and identified several instances where age assignment is erroneous (Tables S2, S4). These include Sillon X+Dan at Sidi Daoui, Morocco (locality #1155), which is assigned to the Maastrichtian-Danian whereas it is well known that this horizon corresponds to the mixing by bioturbation of Maastrichtian and Danian layers (Cappetta 1987; Noubhani & Cappetta 1997). Other localities including reworked Cretaceous taxa in Paleogene horizons (Dutheil, 1996; Smith 1999) or occurrences of unverified ages (e.g., van Baal et al. 2013) were also identified (Tables S2, S4).

We compared the species list of K/Pg victims from Gardiner et al.'s dataset (last updated in 2020) with the species having their last occurrence in the Maastrichtian from Guinot & Condamine (2023) (last updated 2022). Applying the filters of Gardiner et al. (no occurrence with age ranges >15 myr, no species published after 2020), we identified 14 species missing in Gardiner et al. (Table S5), and a large number of key occurrences, including the type locality of obligate Maastrichtian or Paleogene species (e.g., *Squalicorax pristodontus, Pseudocorax* *affinis*), which correspond to ancient literature (e.g., Agassiz 1833-44). We did not check other species (e.g., ptychodontid species).

***Calculation of extinction magnitude***

Deep-learning models provide predictions of diversity trajectories through time by introducing unobserved lineages that lack taxonomic identity, phylogenetic continuity, or morphological definition. These inferred entities cannot be independently validated or falsified and therefore do not constitute biological lineages. This approach and alternatives focus on predicted species richness, thus overlooking underlying speciation and extinction rates, except for the K/Pg event. We argue that the simple calculation of the "*magnitude of species loss*" based on absolute inferred species richness values in Gardiner et al. is irrelevant. This calculation of net change (%) in species richness between two time intervals simply compares absolute richness regardless of the number of apparitions/extinctions creating the observed difference, and its use is not supported by any references in Gardiner et al.

We calculated extinction magnitude (%) following the conventional method e×100/D, where *e* is the number of taxa going extinct and D is the standing diversity (Sepkoski 1984; Foote 2000; Stanley 2016), i.e. the richness at the start of the time bin. We first calculated this percentage of extinction on the raw data from Gardiner et al. (i.e., with dubious occurrences and outlier occurrences creating a high proportion of extinction survivors) for species. We then calculated the extinction percentage once species erroneously considered as survivors in Gardiner et al.'s dataset were treated as typical Cretaceous or Paleogene species (Table S6). We repeated the same procedure for genus-level data (Table S7). We also calculated similar estimates of extinction magnitude but using D as the Total Richness (all taxa present in the time bin, including singletons and taxa originating in the bin) for comparison purposes with other studies (e.g., Bambach 2006).

Using the raw fossil-occurrence data provided in Gardiner et al., we find an extinction magnitude of 60.55 (e=66, D=109) to 64.32% (e=146, D=227) for species and 30.83 (e=37, D=120) to 33.77% (e=51, D=151) for genera, while Gardiner et al.'s calculation found a change in taxonomic richness of −32% ±2% for species and −23% ±1% for genera. Once dubious occurrences and taxonomy were corrected, the magnitude of the K/Pg extinction reaches between 90.29% (e=93, D=103) and 93.53% (e=188, D=201) for species and 63.73% (e=65, D=102) to 65.29% (e=79, D=121) for genera using the conventional method.

### Supplemental Figures

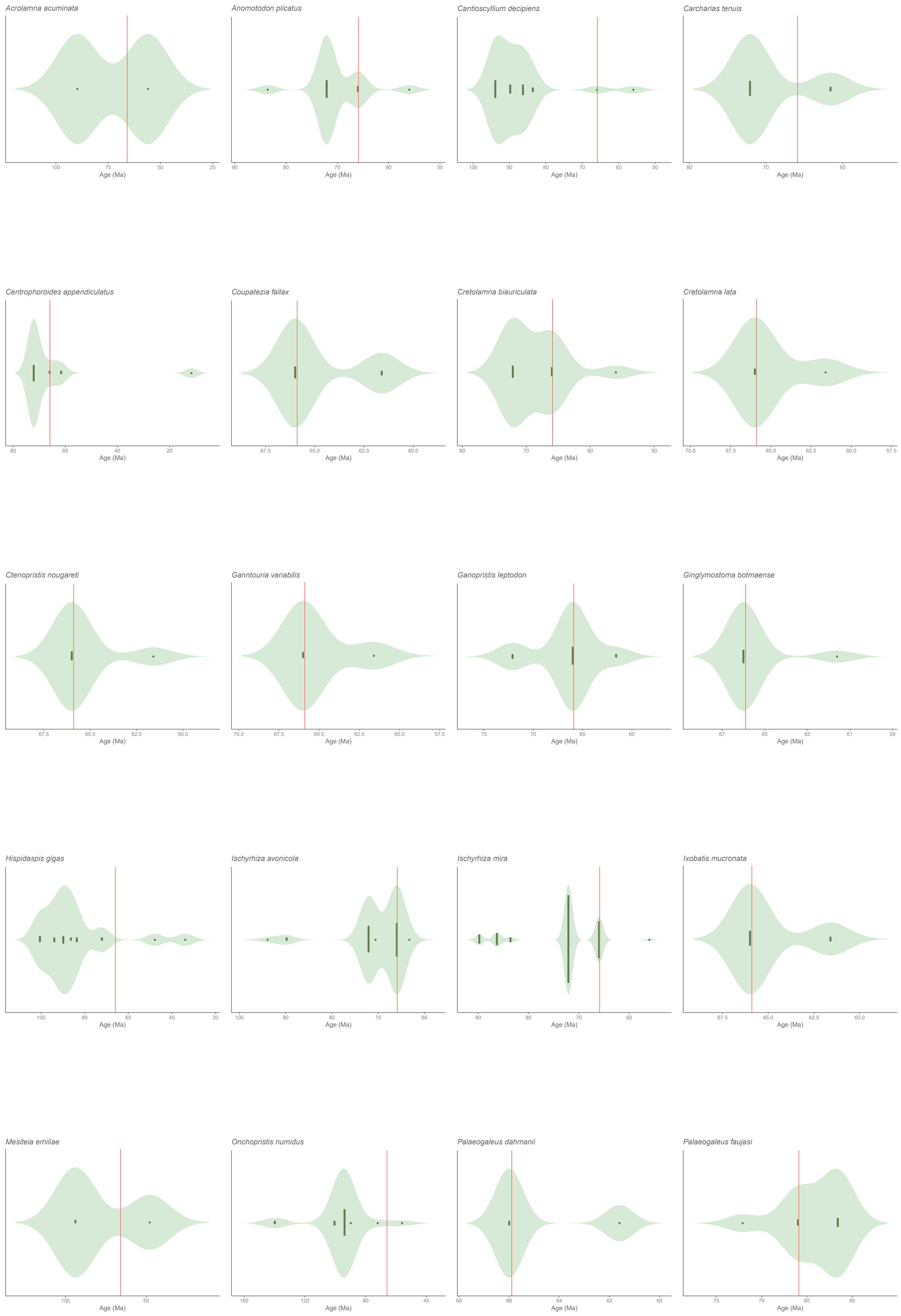


*Figure S1 continued next page*

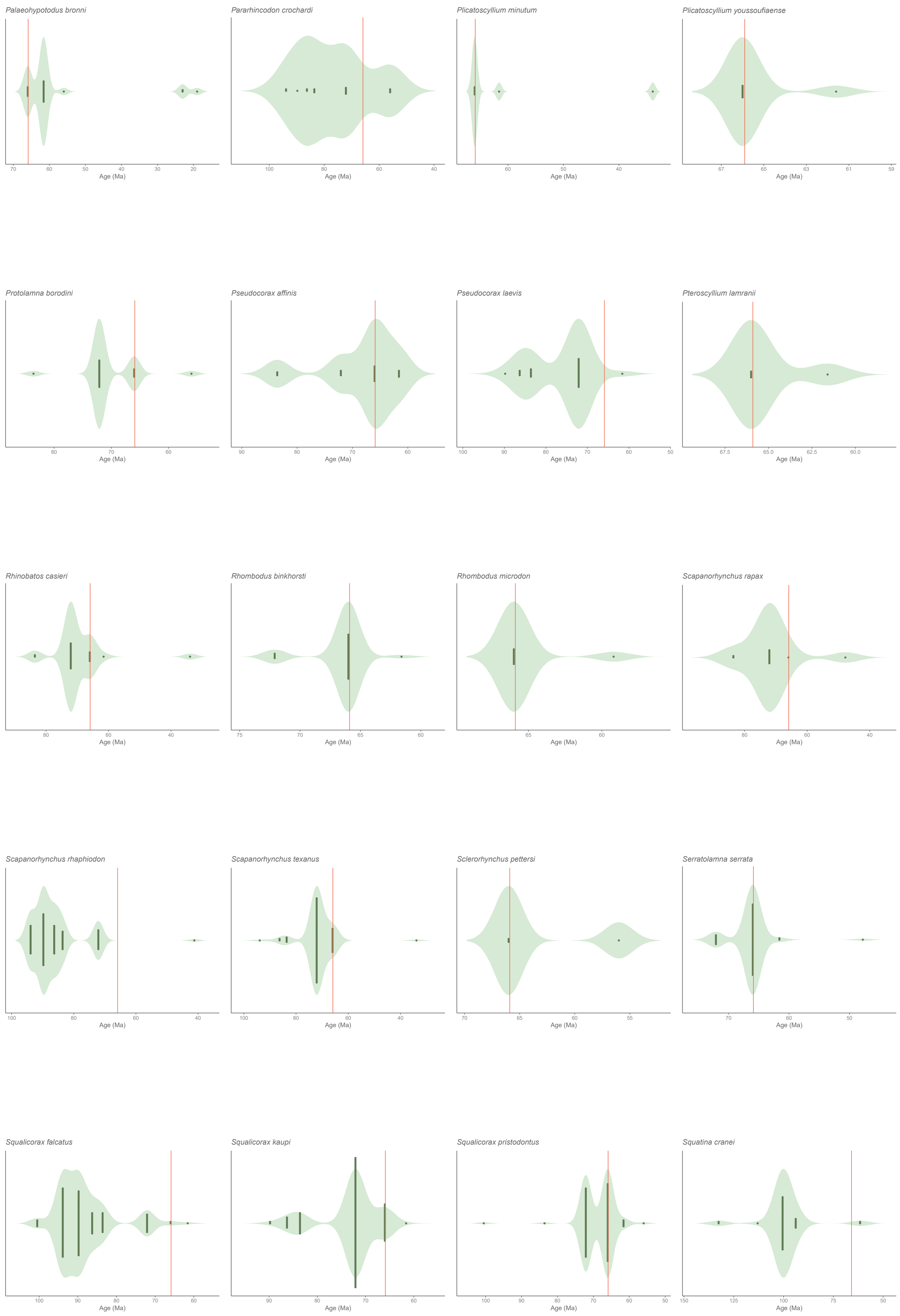


*Figure S1 continued next page*

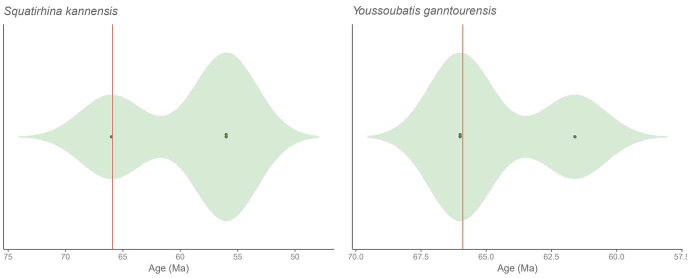


**Figure S1.** Violin plots representing the temporal distribution of occurrences of typical Cretaceous species. The K/Pg boundary is represented by the red vertical bar. Note that occurrences at the oldest end of the distribution were not checked.

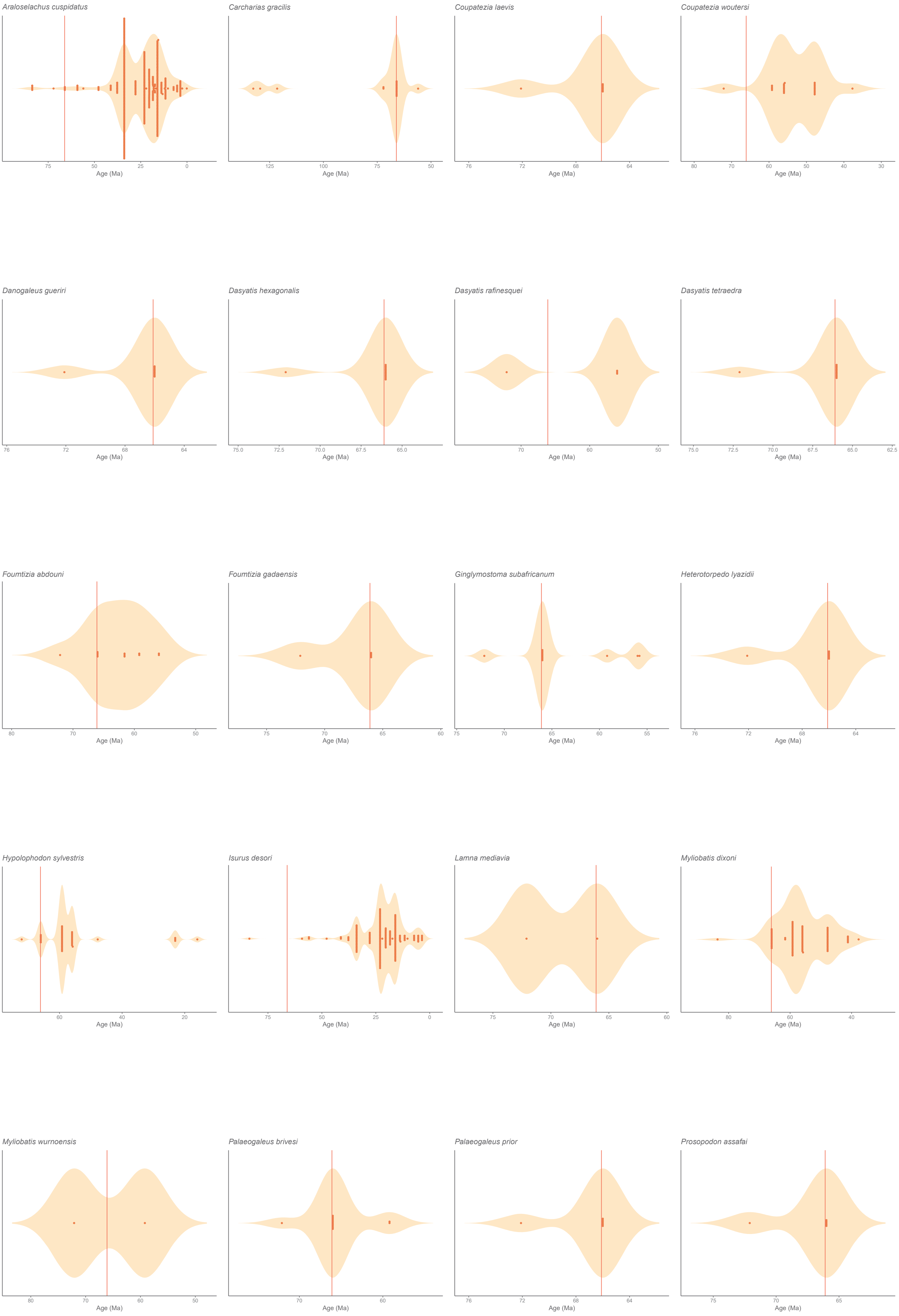


*Figure S2 continued next page*

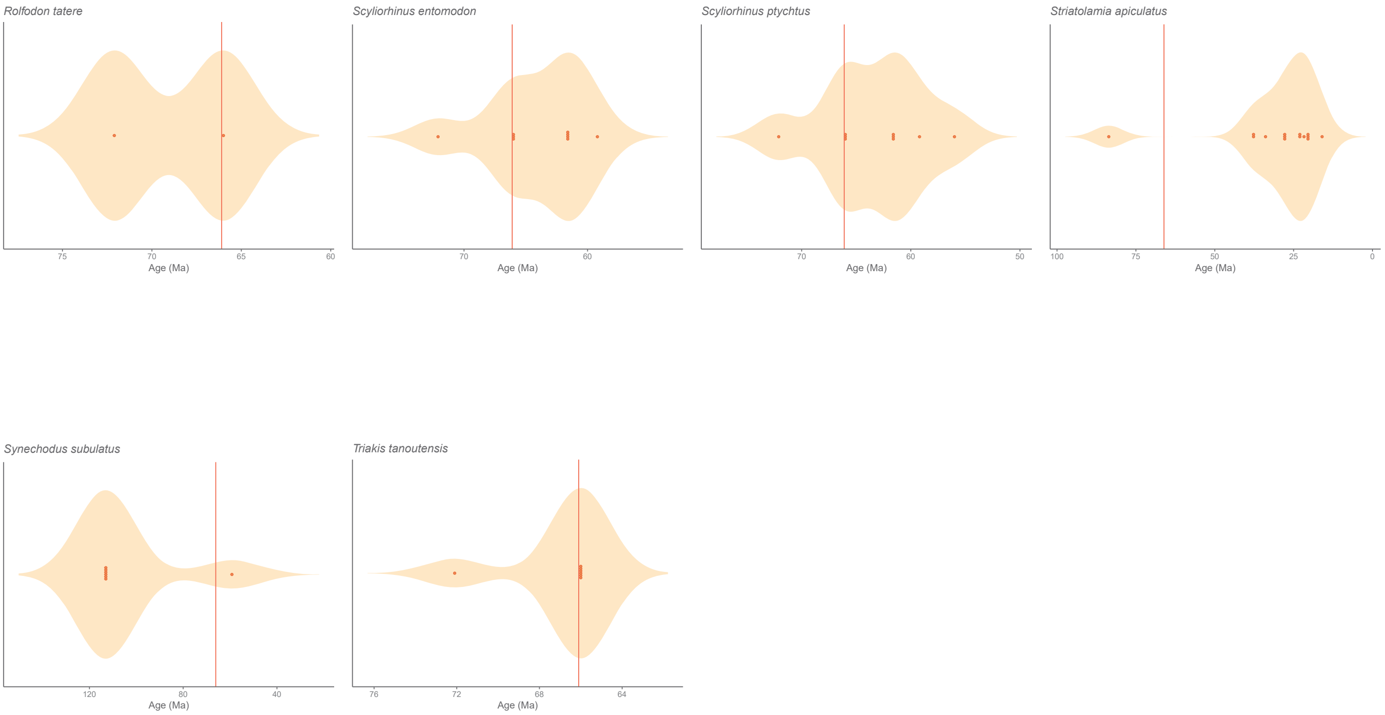


**Figure S2.** Violin plots representing the temporal distribution of occurrences of typical Paleogene species. The K/Pg boundary is represented by the red vertical bar. Note that occurrences at the youngest end of the distribution were not checked.

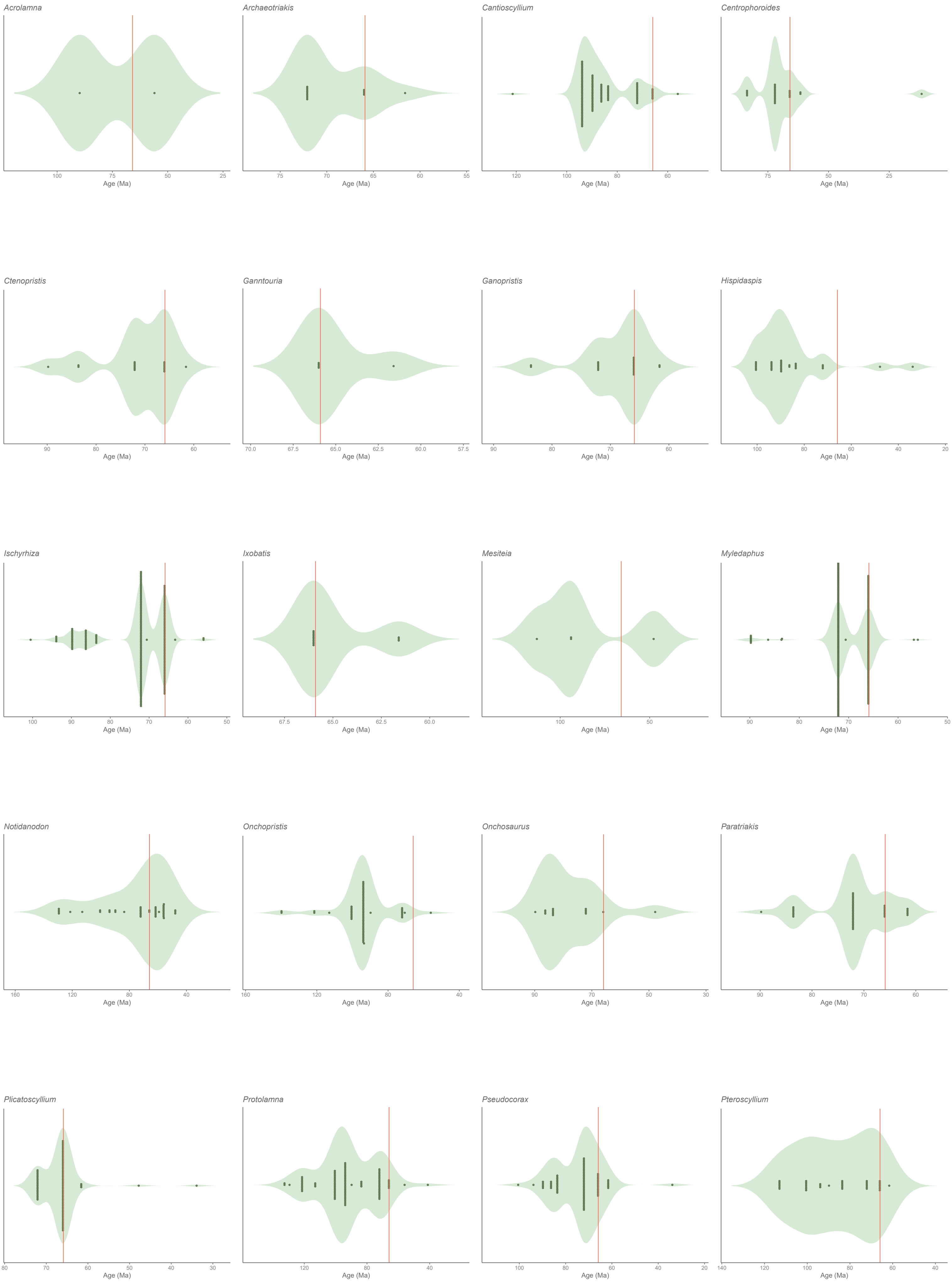


*Figure S3 continued next page*

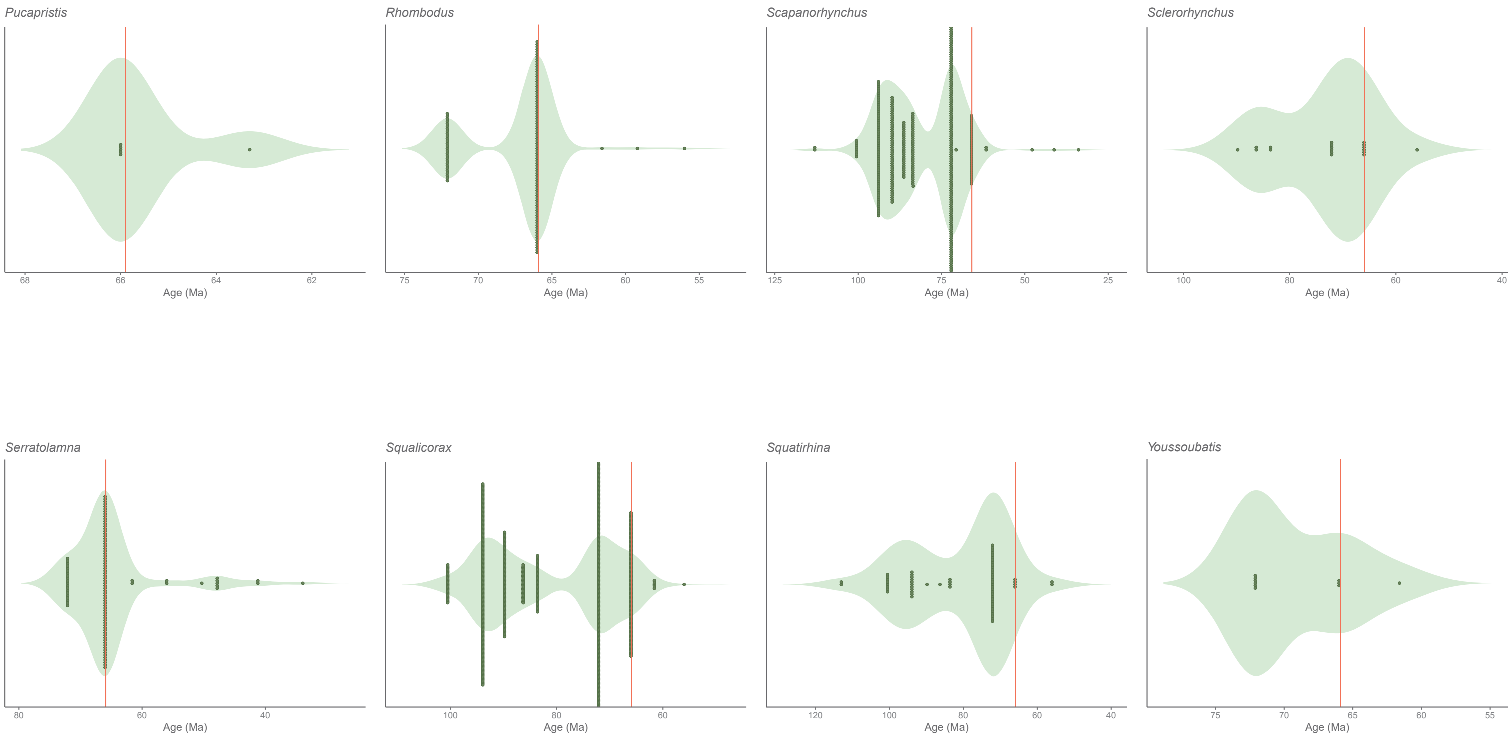


**Figure S3.** Violin plots representing the temporal distribution of occurrences of typical Cretaceous genera. The K/Pg boundary is represented by the red vertical bar. Note that occurrences at the oldest end of the distribution were not checked.

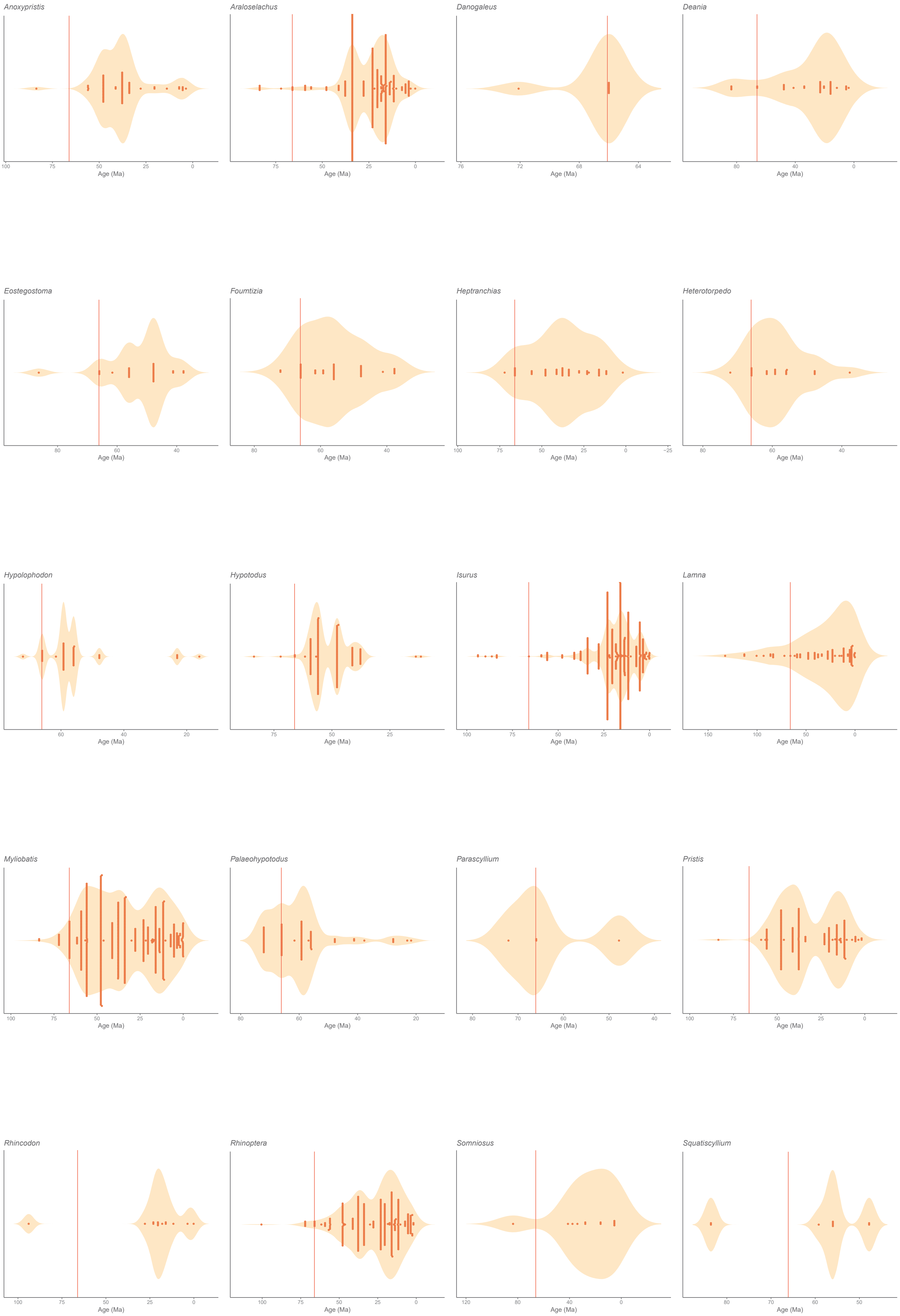


*Figure S4 continued next page*

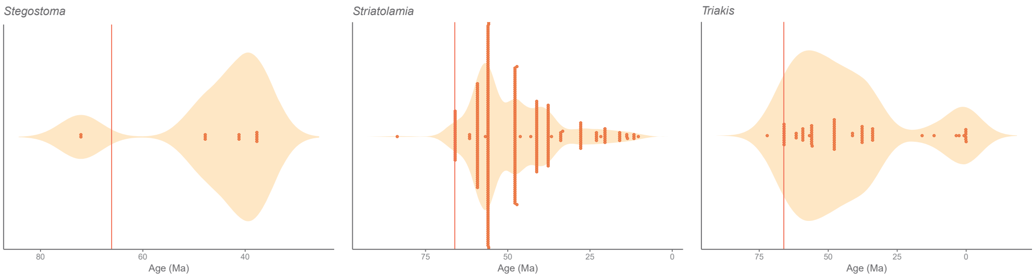


**Figure S4.** Violin plots representing the temporal distribution of occurrences of typical Paleogene genera. The K/Pg boundary is represented by the red vertical bar. Note that occurrences at the youngest end of the distribution were not checked.

#### Supplemental Tables

**Table S1.** Species identified as K/Pg victims (minimum age of youngest occurrence at 66 Ma) based on Data S1 of Gardiner et al. (2026)

| | accepted_name | Age Max | Age Min |
|---|---|---|---|
| 1 | Ankistrorhynchus major | 72.1 | 66 |
| 2 | Anomotodon laevis | 72.1 | 66 |
| 3 | Archaeolamna kopingensis | 113 | 66 |
| 4 | Archaeotriakis rochelleae | 83.6 | 66 |
| 5 | Ataktobatis variabilis | 72.1 | 66 |
| 6 | Biropristis landbecki | 83.6 | 66 |
| 7 | Brachaelurus hornerstownensis | 72.1 | 66 |
| 8 | Brachyrhizodus wichitaensis | 86.3 | 66 |
| 9 | Britobatos primarmata | 86.3 | 66 |
| 10 | Cantioscyllium estesi | 100.5 | 66 |
| 11 | Cantioscyllium meyeri | 83.6 | 66 |
| 12 | Carcharias hardingi | 83.6 | 66 |
| 13 | Carcharias holmdelensis | 86.3 | 66 |
| 14 | Carcharias samhammeri | 86.3 | 66 |
| 15 | Cenocarcharias tenuiplicatus | 100.5 | 66 |
| 16 | Centroscymnus schmidi | 72.1 | 66 |
| 17 | Chiloscyllium gaemersi | 83.6 | 66 |
| 18 | Chiloscyllium greeni | 121.4 | 66 |
| 19 | Coupatezia ambroggii | 72.1 | 66 |
| 20 | Coupatezia elevata | 72.1 | 66 |
| 21 | Coupatezia reniformis | 72.1 | 66 |
| 22 | Coupatezia trempina | 72.1 | 66 |
| 23 | Coupatezia turneri | 72.1 | 66 |
| 24 | Cretalamna feldmanni | 72.1 | 66 |
| 25 | Cretolamna nigeriana | 72.1 | 66 |
| 26 | Cretorectolobus olsoni | 83.6 | 66 |
| 27 | Cretoxyrhina mantelli | 100.5 | 66 |
| 28 | Dalpiazia stromeri | 83.5 | 66 |
| 29 | Dasyatis branisai | 72.1 | 66 |
| 30 | Dasyatis commercensis | 83.6 | 66 |
| 31 | Dasyatis martini | 72.1 | 66 |
| 32 | Dasyatis molinoensis | 72.1 | 66 |
| 33 | Dasyatis newegyptensis | 72.1 | 66 |
| 34 | Dasyatis northdakotaensis | 72.1 | 66 |
| 35 | Dasyatis schaefferi | 72.1 | 66 |
| 36 | Dasyrhombodus bondoni | 72.1 | 66 |
| 37 | Echinorhinus maremagnum | 72.1 | 66 |
| 38 | Eoetmopterus supracretaceus | 83.6 | 66 |
| 39 | Eostriatolamia subulata | 113 | 66 |
| 40 | Galagadon nordquistae | 72.1 | 66 |
| 41 | Galeorhinus girardoti | 83.6 | 66 |
| 42 | Gibbechinorhinus lewyi | 72.1 | 66 |
| 43 | Ginglymostoma cuspidata | 72.1 | 66 |
| 44 | Ginglymostoma erramii | 72.1 | 66 |
| 45 | Hamrabatis weltoni | 72.1 | 66 |
| 46 | Harranahynchus minutadens | 72.1 | 66 |
| 47 | Hemiscyllum hermani | 72.1 | 66 |
| 48 | Heterodontus granti | 83.6 | 66 |
| 49 | Hypsobatis weileri | 72.1 | 66 |
| 50 | Igdabatis indicus | 83.6 | 66 |
| 51 | Igdabatis marmii | 72.1 | 66 |
| 52 | Igdabatis sigmodon | 72.1 | 66 |
| 53 | Ischyrhiza chilensis | 72.1 | 66 |
| 54 | Ischyrhiza hartenbergeri | 72.1 | 66 |
| 55 | Ischyrhiza nigeriensis | 72.1 | 66 |
| 56 | Kiestus texana | 100.5 | 66 |
| 57 | Microetmopterus wardi | 72.1 | 66 |
| 58 | Myledaphus araucanus | 72.1 | 66 |
| 59 | Myledaphus bipartitus | 83.6 | 66 |
| 60 | Myledaphus pustulosus | 72.1 | 66 |
| 61 | Myliobatis foxhillsensis | 72.1 | 66 |
| 62 | Myliobatis leidyi | 72.1 | 66 |
| 63 | Odontaspis aculeatus | 83.6 | 66 |
| 64 | Onchosaurus pharao | 93.3 | 66 |
| 65 | Palaeogaleus navarroensis | 72.1 | 66 |
| 66 | Paraginglymostoma bloti | 72.1 | 66 |
| 67 | Paranomotodon angustidens | 113 | 66 |
| 68 | Paranomotodon toddi | 86.3 | 66 |
| 69 | Paraorthacodus andersoni | 83.6 | 66 |
| 70 | Paraorthacodus conicus | 83.6 | 66 |
| 71 | Parapalaeobates atlanticus | 83.5 | 66 |
| 72 | Parasquatina zitteli | 72.1 | 66 |
| 73 | Paratrygonorrhina amblysoda | 72.1 | 66 |
| 74 | Plicatoscyllium antiquum | 72.1 | 66 |
| 75 | Plicatoscyllium derameei | 72.1 | 66 |
| 76 | Plicatoscyllium gharbii | 72.1 | 66 |
| 77 | Plicatoscyllium globidens | 86.3 | 66 |
| 78 | Plicatoscyllium lehneri | 72.1 | 66 |
| 79 | Plicatoscyllium pectinatum | 72.1 | 66 |
| 80 | Proetmopterus hemmooriensis | 72.1 | 66 |
| 81 | Proheterodontus creamridgensis | 72.1 | 66 |
| 82 | Protoplatyrhina renae | 93.9 | 66 |
| 83 | Pseudocorax granti | 93.9 | 66 |
| 84 | Pseudodontaspis herbsti | 83.6 | 66 |
| 85 | Pseudoginglymostoma erguitaense | 72.1 | 66 |
| 86 | Pseudoginglymostoma idiri | 72.1 | 66 |
| 87 | Pseudohypolophus mcnultyi | 129.4 | 66 |
| 88 | Ptychodus atcoensis | 89.8 | 66 |
| 89 | Ptychodus cyclodontis | 72.1 | 66 |
| 90 | Ptychodus decurrens | 113 | 66 |
| 91 | Ptychodus mortoni | 93.9 | 66 |
| 92 | Ptychotrygon agujaensis | 83.6 | 66 |
| 93 | Ptychotrygon clementsi | 72.1 | 66 |

**Table S1 (continued)**

| | accepted_name | Age Max | Age Min |
|---|---|---|---|
| 94 | Ptychotrygon cuspidata | 83.6 | 66 |
| 95 | Ptychotrygon slaughteri | 100.5 | 66 |
| 96 | Ptychotrygon texana | 72.1 | 66 |
| 97 | Ptychotrygon triangularis | 100.5 | 66 |
| 98 | Ptychotrygon vermiculata | 93.9 | 66 |
| 99 | Ptychotrygon winni | 72.1 | 66 |
| 100 | Pucabatis hoffstetteri | 83.6 | 66 |
| 101 | Pucapristis branisi | 72.1 | 66 |
| 102 | Raja farishi | 72.1 | 66 |
| 103 | Raja sudhakari | 72.1 | 66 |
| 104 | Restesia americana | 83.6 | 66 |
| 105 | Rhinobatos craddocki | 72.1 | 66 |
| 106 | Rhinobatos echavei | 83.6 | 66 |
| 107 | Rhinobatos ibericus | 72.1 | 66 |
| 108 | Rhinobatos mariannae | 72.1 | 66 |
| 109 | Rhinobatos uvulatus | 72.1 | 66 |
| 110 | Rhombodus andriesi | 72.1 | 66 |
| 111 | Rhombodus ibericus | 72.1 | 66 |
| 112 | Rhombodus levis | 86.3 | 66 |
| 113 | Rhombodus meridionalis | 83.5 | 66 |
| 114 | Schizorhiza stromeri | 83.6 | 66 |
| 115 | Sclerorhynchus atavus | 86.3 | 66 |
| 116 | Scyliorhinus ivagrantae | 72.1 | 66 |
| 117 | Scyliorhinus luypaertsi | 72.1 | 66 |
| 118 | Scyliorhinus moosi | 72.1 | 66 |
| 119 | Serratolamna africana | 72.1 | 66 |
| 120 | Serratolamna caraibaea | 83.6 | 66 |
| 121 | Serratolamna khderii | 83.6 | 66 |
| 122 | Serratolamna maroccana | 83.6 | 66 |
| 123 | Sphenodus longidens | 100.5 | 66 |
| 124 | Squalicorax africanus | 83.5 | 66 |
| 125 | Squalicorax bassanii | 83.6 | 66 |
| 126 | Squalicorax benguerirensis | 72.1 | 66 |
| 127 | Squalicorax kugleri | 72.1 | 66 |
| 128 | Squalicorax lindstromi | 100.5 | 66 |
| 129 | Squalicorax microserratus | 83.6 | 66 |
| 130 | Squalus argentinensis | 72.1 | 66 |
| 131 | Squalus ballingsloevensis | 72.1 | 66 |
| 132 | Squalus balsvikensis | 72.1 | 66 |
| 133 | Squalus huntensis | 72.1 | 66 |
| 134 | Squalus vondermarcki | 83.6 | 66 |
| 135 | Squatigaleus atlasi | 83.6 | 66 |
| 136 | Squatigaleus sulphurensis | 72.1 | 66 |
| 137 | Squatina hassei | 86.3 | 66 |
| 138 | Synechodus lerichei | 86.3 | 66 |
| 139 | Synechodus turneri | 83.6 | 66 |
| 140 | Tanoutia iminensis | 72.1 | 66 |
| 141 | Texabatis corrugata | 72.1 | 66 |
| 142 | Texatrygon hooveri | 100.5 | 66 |
| 143 | Tomewingia problematica | 72.1 | 66 |
| 144 | Vascobatis albaitensis | 72.1 | 66 |
| 145 | Walteraja exigua | 72.1 | 66 |
| 146 | Xampylodon dentatus | 86.3 | 66 |

**Table S2.** Survivor species identified as such from Dataset S1 of Gardiner et al. (2026) and accompanying issues and corrections. Species names highlighted in green indicate typical Cretaceous species with corresponding problematic Cenozoic occurrences. Species names highlighted in orange indicate typical Paleogene species with corresponding problematic Cretaceous occurrences. Species names in red font highlight invalid or dubious taxon names.

| accepted_name | Age Max | Age Min | Outlier occurrences | Age issue | Occ._Nber | Occ._Ref | evidence | Taxonomic issue |
|---|---|---|---|---|---|---|---|---|
| *Acrolamna acuminata* | 93.9 | 56 | 1 Cenozoic occ. | | PBDB_23491 | Cook and Ramsdell (1991) | no_evidence | |
| *Anomotodon plicatus* | 86.3 | 56 | 1 Cenozoic occ. | | L_28636 | Reinecke & Engelhard 1997 | with_evidence | Open nomenclature: as *Anomotodon plicatus* ? in corresponding Paleogene reference |
| *Araloselachus cuspidatus* | 83.6 | 0.0117 | 5 Cretaceous occ. | | PBDB_23374; PBDB_12886; PBDB_12889; PBDB_12890; PBDB_12892 | Gallagher 1984; Berry 1916 | no_evidence | |
| *Cantioscyllium decipiens* | 100.5 | 56 | 1 Cenozoic occ. | Reworked/mixed Maastrichtian & Danian | L_21081 | Smith et al. 1999 | with_evidence | |
| *Carcharias gracilis* | 132.6 | 41.2 | 5 Cretaceous occ. | | L_28840; PBDB_16300; L_5414; L_5409; L_5403 | Pereda-Suberbiola et al. 2015; Cappetta and Corral 1999; Biddle 1993b | both | Open nomenclature: the 2 occ. "with_evidence" are *Carcharias* aff. *gracilis* |
| *Carcharias tenuis* | 83.6 | 61.6 | 3 Cenozoic occ. | | L_1534; PBDB_103; PBDB_10306 | Siverson 1993b; Siverson 1995 | both | Open nomenclature: the 3 occ. are as *Carcharias* aff. *tenuis* |
| *Centrophoroides appendiculatus* | 83.6 | 11.63 | 3 Cenozoic occ. | | PBDB_25406; L_4309; L_5490 | Cope 1867; van Baal et al. 2013; Dutheil 1996 | both | The occ. van Baal et al. is not based on evidence |
| *Centroscymnus praecursor* | 83.6 | 61.6 | | | | | | |
| *Coupatezia fallax* | 72.1 | 61.6 | 3 Cenozoic occ. | Reworking (incl. loc. 1155) | PBDB_19943; PBDB_19797; L_551 | Noubhani and Cappetta 1997; Dutheil 1996 | both | The occurrence PBDB_19943 from Youssoufia Level 0 is erroneous, see Noubhani & Cappetta (1997): "les dents de cette espèce ne se rencontrent que dans les niveaux supérieurs du Maastrichtien" |
| *Coupatezia laevis* | 72.1 | 61.6 | 1 Cretaceous occ. | Reworking (loc. 1155) | PBDB_19798 | Noubhani and Cappetta 1997 | with_evidence | |
| *Coupatezia woutersi* | 72.1 | 33.9 | 1 Cretaceous occ. | | L_3668 | Prasad and Sahni 1987 | with_evidence | |

**Table S2 (continued).**

| | | | | | | | | |
|---|---|---|---|---|---|---|---|---|
| *Crassescyliorhinus germanicus* | 86.3 | 61.6 | | | | | | |
| *Cretolamna appendiculata* | 113 | 33.9 | | | | | | Records from the Late Eocene based on a poorly preserved tooth (Vasquez and Pimiento 2014) that does not share characters with the genus (Siversson et al 2013), which is unknown above the Early Eocene |
| *Cretolamna biauriculata* | 86.3 | 56 | 1 Cenozoic occ. | | L_30441 | Da Silva 2007 | with_evidence | |
| *Cretolamna lata* | 72.1 | 61.6 | 1 Cenozoic occ. | | L_2517 | Yarkov & Popov 1998 | no_evidence | |
| *Ctenopristis nougareti* | 72.1 | 61.6 | 1 Cenozoic occ. | | PBDB_10050 | Tong and Meylan 2013 | with_evidence | This occ. is not based on evidence |
| *Danogaleus gueriri* | 72.1 | 61.6 | 1 Cretaceous occ. | Reworking (loc. 1155) | PBDB_19793 | Noubhani and Cappetta 1997 | with_evidence | |
| *Dasyatis hexagonalis* | 72.1 | 59.2 | 1 Cretaceous occ. | Reworking (loc. 1155) | PBDB_19794 | Noubhani and Cappetta 1997 | with_evidence | |
| *Dasyatis rafinesquei* | 72.1 | 47.8 | 1 Cretaceous occ. | | L_29378 | Wroblewski 2004 | with_evidence | Cretaceous occurrence entered as "*Dasyatis* Rafinesque" (*genus name* + author) and not *Dasyatis rafinesquei* |
| *Dasyatis tetraedra* | 72.1 | 61.6 | 1 Cretaceous occ. | Reworking (loc. 1155) | PBDB_19795 | Noubhani and Cappetta 1997 | with_evidence | |
| *Foumtizia abdouni* | 72.1 | 47.8 | 1 Cretaceous occ. | Reworking (loc. 1155) | PBDB_19787 | Noubhani and Cappetta 1997 | with_evidence | |
| *Foumtizia gadaensis* | 72.1 | 61.6 | 1 Cretaceous occ. | Reworking (loc. 1155) | PBDB_19788 | Noubhani and Cappetta 1997 | with_evidence | |
| *Ganntouria variabilis* | 72.1 | 61.6 | 1 Cenozoic occ. | Reworking (loc. 1155) | PBDB_19783 | Noubhani and Cappetta 1997 | with_evidence | |
| *Ganopristis leptodon* | 83.6 | 61.6 | 2 Cenozoic occ. | | L_4300; L_5508 | Dutheil 1996; van Baal et al. 2013 | both | van Baal et al. is considered as "with evidence" whereas no evidence is provided |
| *Ginglymostoma botmaense* | 72.1 | 61.6 | 1 Cenozoic occ. | Reworking (loc. 1155) | PBDB_19781 | Noubhani and Cappetta 1997 | with_evidence | |
| *Ginglymostoma subafricanum* | 72.1 | 47.8 | 1 Cretaceous occ. | Reworking (loc. 1155) | PBDB_19782 | Noubhani and Cappetta 1997 | with_evidence | |
| *Hemiscyllium hermani* | 83.6 | 61.6 | | | | | | |

**Table S2 (continued).**

| | | | | | | | | |
|---|---|---|---|---|---|---|---|---|
| *Heterodontus rugosus* | 100.5 | 56 | | | | | | |
| *Heterotorpedo lyazidii* | 72.1 | 61.6 | 1 Cretaceous occ. | Reworking (loc. 1155) | PBDB_19799 | Noubhani and Cappetta 1997 | with_evidence | |
| *Hexanchus microdon* | 100.5 | 48.6 | | | | | | |
| *Hispidaspis gigas* | 113 | 33.9 | 2 Cenozoic occ. | | L_20698; L_20691 | Li 1997 | with_evidence | |
| *Hypolophodon sylvestris* | 72.1 | 5.333 | 1 Cretaceous occ. | | L_29649 | Bogan and Gallina 2011 | with_evidence | |
| *Ischyrhiza avonicola* | 100.5 | 63.3 | 1 Cenozoic occ. | | PBDB_14116 | Van Valen and Sloan 1965 | no_evidence | Open nomenclature: as "cf. *Ischyrhiza avonicola* " |
| *Ischyrhiza mira* | 93.9 | 56 | 1 Cenozoic occ. | | PBDB_23502 | Cook and Ramsdell 1991 | no_evidence | |
| *Isurus desori* | 83.6 | 2.58 | 1 Cretaceous occ. | | PBDB_12102 | Chapman 1918 | no_evidence | |
| *Ixobatis mucronata* | 72.1 | 61.6 | 3 Cenozoic occ. | Reworking (loc. 1155) | PBDB_13713; PBDB_19778; PBDB_13713 | Hirayama and Tong 2003; Noubhani and Cappetta 1995; Tong and Meylan 2013 | with_evidence | 2 occ. doubled |
| Lamna mediavia | 72.1 | 56 | 1 Cretaceous occ. | | PBDB_10082 | Leriche 1942 | no_evidence | *Lamna mediavia* Leriche, 1942 was described from the Danian (Midway Fm.) of Prairie Creek, Wilcox Co., Alabama, U.S.A. Was cited earlier by Leriche (1940, p. 590), but without illustration. Considered as a sub-species of *Otodus minor* Leriche, 1909 by Zhelezko *in* Zhelezko & Kozlov (1999). Also included in *Otodus* by Cappetta (2006). |
| *Mesiteia emiliae* | 100.5 | 47.8 | 1 Cenozoic occ. | | PBDB_10599 | Carnevale et al. 2014 | no_evidence | This occurrence is considered dubious as indicated in the reference |
| *Myliobatis dixoni* | 83.6 | 33.9 | 1 Cretaceous occ. | | PBDB_12894 | Berry 1916 | no_evidence | As *Myliobatis obesus* |
| *Myliobatis wurnoensis* | 72.1 | 56 | 1 Cretaceous occ. | Dubious stratigraphic origins | PBDB_12328 | O'Leary et al. 2019 | with_evidence | |

**Table S2 (continued).**

| | | | | | | | | |
|---|---|---|---|---|---|---|---|---|
| *Onchopristis numidus* | 145 | 56 | 1 Cenozoic occ. | Age of occurrence indicated as "Possibly Late Maastrichtian-Paleocene" and "Currently we have poor field records regarding the stratigraphic layer in which this taxon was collected." in O'Leary et al (2019; p. 94) | PBDB_17257 | O'Leary et al. 2019 | with_evidence | |
| *Palaeogaleus brivesi* | 72.1 | 56 | 1 Cretaceous occ. | Reworking (loc. 1155) | PBDB_19789 | Noubhani and Cappetta 1997 | with_evidence | |
| *Palaeogaleus dahmanii* | 72.1 | 61.6 | 1 Cenozoic occ. | Reworking (loc. 1155) | PBDB_19790 | Noubhani and Cappetta 1997; Dutheil 1996 | with_evidence | |
| *Palaeogaleus faujasi* | 83.6 | 61.6 | 6 occ. younger | | L_1508; L_1515; L_155; L_423; L_4238; L_5503 | Adolfssen and Ward 2015; Nilsson 2003; Dutheil 1996 | both | Open nomenclature: the 5 occ. with evidence are as *P.* aff. *faujasi* , the other is without evidence (Dutheil 1996) |
| *Palaeogaleus prior* | 72.1 | 61.6 | 1 Cretaceous occ. | Reworking (loc. 1155) | PBDB_19791 | Noubhani and Cappetta 1997 | with_evidence | |
| *Palaeohypotodus bronni* | 72.1 | 19 | 9 Cenozoic occ. | | L_1503; L_1511; L_4224; L_4234; L_4280; PBDB_23524; PBDB_24533; PBDB_24681; PBDB_17409 | Adolfssen and Ward 2015; Adolfssen 2012; van Baal et al. 2013; Jagt 1996; Chapman 1918; Chapman and Pritchard 1904; Chapman 1918 | both | Open nomenclature: as cf. or aff. for 4 Cretaceous occurrences, others are without evidence |
| *Pararhincodon crochardi* | 100.5 | 56 | 3 Cenozoic occ. | Reworking (Smith et al. 1999) | L_21014; L_21046; L_21547 | Smith et al. 1999; Moreau & Mathis 2000 | with_evidence | Open nomenclature: reported as *Pararhincodon* cf. *corchardi* in Moreau & Mathis (2000) but not indicated in GEA's dataset |
| *Pararhincodon groessensi* | 83.6 | 61.6 | | | | | | |
| *Parasquatina cappettai* | 72.1 | 61.6 | | | | | | |
| *Paratriakis curtirostris* | 86.3 | 61.6 | | | | | | Belongs to another genus (*Palaeotriakis* ) (see Guinot et al. 2013) |

**Table S2 (continued).**

| | | | | | | | | |
|---|---|---|---|---|---|---|---|---|
| *Plicatoscyllium minutum* | 72.1 | 33.9 | 2 Cenozoic occ. | Reworked (Dutheil 1996) | L_20696; L_5494 | Li 1997; Dutheil 1996 | both | |
| *Plicatoscyllium youssoufiaense* | 72.1 | 61.6 | 1 Cenozoic occ. | Reworking (loc. 1155) | PBDB_19780 | Noubhani and Cappetta 1997 | with_evidence | |
| *Prosopodon assafai* | 72.1 | 61.6 | 1 Cretaceous occ. | Reworking (loc. 1155) | PBDB_19777 | Noubhani and Cappetta 1997 | with_evidence | Name *Prosopodon* replaced by *Phosphatodon* (see Cappetta 2012) |
| *Protolamna borodini* | 86.3 | 56 | 1 Cenozoic occ. | | L_28638 | Reinecke & Engelhard 1997 | with_evidence | Open nomenclature: as *Cretodus borodini* ? in original publication |
| *Pseudocorax affinis* | 86.3 | 61.6 | 1 Cenozoic occ. | | L_2516 | Yarkov & Popov 1998 | no_evidence | |
| *Pseudocorax laevis* | 93.9 | 61.6 | 1 Cenozoic occ. | Reworked | L_55 | Dutheil 1996 | no_evidence | |
| *Pteroscyllium lamranii* | 72.1 | 61.6 | 1 Cenozoic occ. | Reworking (loc. 1155) | PBDB_19784 | Noubhani and Cappetta 1997 | with_evidence | |
| *Rhinobatos casieri* | 89.8 | 33.9 | 2 Cenozoic occ. | | PBDB_26939; L_5505 | Case 1981; Dutheil 1996 | no_evidence | |
| *Rhombodus binkhorsti* | 83.6 | 61.6 | 1 Cenozoic occ. | Reworked | L_5512 | Dutheil 1996 | no_evidence | |
| *Rhombodus microdon* | 72.1 | 59.2 | 1 Cenozoic occ. | | PBDB_13039 | Solé et al. 2009 | with_evidence | |
| *Rolfodon tatere* | 72.1 | 61.6 | 1 Cretaceous occ. | | PBDB_17638 | Blokland et al. 2019 | with_evidence | |
| *Scapanorhynchus rapax* | 86.3 | 47.8 | 1 Cenozoic occ. | | PBDB_24266 | Abbass 1972 | no_evidence | |
| *Scapanorhynchus rhaphiodon* | 100.5 | 41.2 | 1 Cenozoic occ. | | PBDB_27037 | Arambourg and Joleaud 1943 | no_evidence | |
| *Scapanorhynchus texanus* | 100.5 | 33.9 | 1 Cenozoic occ. | | PBDB_24487 | White 1956 | no_evidence | |
| *Sclerorhynchus pettersi* | 72.1 | 56 | 1 Cenozoic occ. | Reworking | L_2102 | Smith 1999 | with_evidence | Open nomenclature: as *Sclerorhynchus* cf. *pettersi* |
| *Scyliorhinus biddlei* | 72.1 | 61.6 | | | | | | |
| *Scyliorhinus elongatus* | 132.6 | 56 | | | | | | |
| *Scyliorhinus entomodon* | 72.1 | 56 | 1 Cretaceous occ. | Reworking (loc. 1155) | PBDB_19785 | Noubhani and Cappetta 1997 | with_evidence | |
| *Scyliorhinus ptychtus* | 72.1 | 47.8 | 1 Cretaceous occ. | Reworking (loc. 1155) | PBDB_19786 | Noubhani and Cappetta 1997 | with_evidence | Open nomenclature: as *Scyliorhinus* aff. *ptychtus* |
| *Serratolamna serrata* | 86.3 | 47.8 | 3 Cenozoic occ. | Reworking (Dutheil 1996) | PBDB_24267; L_4281; L_5499 | Abbass 1972; van Baal et al. 2013; Dutheil 1996 | both | van Baal et al. is considered as "with evidence" whereas no evidence is provided |
| *Sphenodus lundgreni* | 89.8 | 55.8 | | | | | | |
| *Squalicorax falcatus* | 113 | 61.6 | 1 Cenozoic occ. | | L_2274 | Shourd & Winter 1980 | with_evidence | |
| *Squalicorax kaupi* | 93.9 | 61.6 | 1 Cenozoic occ. | Reworking | L_550 | Dutheil 1996 | no_evidence | |

**Table S2 (continued).**

| | | | | | | | | |
|---|---|---|---|---|---|---|---|---|
| *Squalicorax pristodontus* | 113 | 56 | 6 Cenozoic occ. | | L_3922; L_2515; L_4285; L_430; L_4302; L_4306 | Da Silva 2007; Yarkov & Popov 1998; van Baal et al. 2013 | both | van Baal et al. is considered as "with evidence" whereas no evidence is provided |
| *Squalus gabrielsoni* | 72.1 | 61.6 | | | | | | |
| *Squatina cranei* | 139.8 | 61.6 | 2 occ. younger | | L_4219; L_4246 | Adolfssen 2012 | with_evidence | These occ. are from an unpublished thesis (Adolfssen 2012), and were published as *Squatina* sp. Adolfssen & Ward (2014, 2015) |
| *Squatirhina kannensis* | 72.1 | 56 | 2 Cenozoic occ. | Reworked | L_21049; L_21078 | Smith et al. 1999 | with_evidence | Open nomenclature: as *Squatirhina* cf. *kannensis* |
| *Striatolamia apiculatus* | 83.6 | 11.63 | 1 Cretaceous occ. | | PBDB_12099 | Chapman 1918 | no_evidence | Junior synonym (pars.) of Cenozoic *Cosmopolitodus hastalis* (Agassiz, 1843), see Cappetta (2006) |
| *Synechodus subulatus* | 113 | 47.8 | | | | | | Mix between two distinct species: the Albian *Synechodus subulatus* (Rogovich, 1861 *non* Leriche, 1951) (see Sokolskyi & Guinot, 2021) and the Danian *S. subulatus* Leriche, 1951 *non* Rogovich, 1860 (see Hovestadt & Steurbaut, 2023) |
| *Triakis tanoutensis* | 72.1 | 61.6 | 1 Cretaceous occ. | Reworking (loc. 1155) | PBDB_19792 | Noubhani and Cappetta 1997 | with_evidence | |
| *Youssoubatis ganntourensis* | 72.1 | 61.6 | 1 Cenozoic occ. | | PBDB_10051 | Tong and Meylan 2013 | with_evidence | The occ. is not based on evidence |

**Table S3.** Genera identified as K/Pg victims (minimum age of youngest occurrence at 66 Ma) based on Data S1 of Gardiner et al. (2026)

| | genus | Max_Age | Min_Age |
|---|---|---|---|
| 1 | Acanthoscyllium | 86.3 | 66 |
| 2 | Ankistrorhynchus | 89.8 | 66 |
| 3 | Annea | 113 | 66 |
| 4 | Archaeolamna | 113 | 66 |
| 5 | Ataktobatis | 83.5 | 66 |
| 6 | Biropristis | 83.6 | 66 |
| 7 | Brachyrhizodus | 129.4 | 66 |
| 8 | Britobatos | 86.3 | 66 |
| 9 | Cederstroemia | 113 | 66 |
| 10 | Cenocarcharias | 100.5 | 66 |
| 11 | Cretomanta | 100.5 | 66 |
| 12 | Cretorectolobus | 132.6 | 66 |
| 13 | Cretoxyrhina | 113 | 66 |
| 14 | Dalpiazia | 83.5 | 66 |
| 15 | Dasyrhombodus | 72.1 | 66 |
| 16 | Eoetmopterus | 100.5 | 66 |
| 17 | Eostriatolamia | 113 | 66 |
| 18 | Galagadon | 72.1 | 66 |
| 19 | Gibbechinorhinus | 72.1 | 66 |
| 20 | Hamrabatis | 100.5 | 66 |
| 21 | Harranahynchus | 72.1 | 66 |
| 22 | Hypsobatis | 83.5 | 66 |
| 23 | Igdabatis | 83.6 | 66 |
| 24 | Kiestus | 100.5 | 66 |
| 25 | Microetmopterus | 72.1 | 66 |
| 26 | Paraginglymostoma | 121.4 | 66 |
| 27 | Paranomotodon | 113 | 66 |
| 28 | Parapalaeobates | 93.9 | 66 |
| 29 | Paratrygonorrhina | 72.1 | 66 |
| 30 | Peyeria | 100.5 | 66 |
| 31 | Proetmopterus | 72.1 | 66 |
| 32 | Proheterodontus | 72.1 | 66 |
| 33 | Propristiophorus | 72.1 | 66 |
| 34 | Protoplatyrhina | 100.5 | 66 |
| 35 | Protosqualus | 129.4 | 66 |
| 36 | Pseudodontaspis | 83.6 | 66 |
| 37 | Pseudohypolophus | 132.6 | 66 |
| 38 | Pseudoisurus | 113 | 66 |
| 39 | Ptychodus | 113 | 66 |
| 40 | Ptychotrygon | 113 | 66 |
| 41 | Pucabatis | 83.6 | 66 |
| 42 | Restesia | 83.6 | 66 |
| 43 | Schizorhiza | 83.6 | 66 |
| 44 | Squatigaleus | 83.6 | 66 |
| 45 | Tanoutia | 72.1 | 66 |
| 46 | Texabatis | 72.1 | 66 |
| 47 | Texatrygon | 100.5 | 66 |
| 48 | Tomewingia | 72.1 | 66 |
| 49 | Vascobatis | 72.1 | 66 |
| 50 | Walteraja | 72.1 | 66 |
| 51 | Xampylodon | 86.3 | 66 |

**Table S4.** Survivor genera identified as such from Dataset S1 of Gardiner et al. (2026) and accompanying issues and corrections. Genus names highlighted in green indicate typical Cretaceous genera with corresponding problematic Cenozoic occurrences. Genus names highlighted in orange indicate typical Paleogene species with corresponding problematic Cretaceous occurrences. Genus names in red font highlight invalid or dubious taxon names.

| genus | Max_Age | Min_Age | Outlier occurrences | Age issue | Occ._Nber | Occ._Ref | evidence | Taxo issue |
|---|---|---|---|---|---|---|---|---|
| Acrolamna | 93.9 | 56 | 1 Cenozoic occ. | | PBDB_23491 | Cook and Ramsdell (1991) | no_evidence | |
| Anomotodon | 132.6 | 11.63 | | | | | | |
| Anoxypristis | 83.6 | 2.58 | 1 Cretaceous occ. | | PBDB_12925 | Crane (2011) | no_evidence | |
| Araloselachus | 83.6 | 0.0117 | 5 Cretaceous occ. | | PBDB_23374; PBDB_12886; PBDB_12889; PBDB_12890; PBDB_12892 | Berry 1916; Gallagher 1984 | no_evidence | |
| Archaeotriakis | 83.6 | 61.6 | 1 Cenozoic occ. | | L_2035 | Bernardez 1997 | no_evidence | |
| Brachaelurus | 139.8 | 5.333 | | | | | | |
| Cantioscyllium | 129.4 | 56 | 1 Cenozoic occ. | Reworked | L_21081 | Smith et al. 1999 | with_evidence | |
| Carcharias | 132.6 | 0.0117 | | | | | | |
| Centrophoroides | 89.8 | 11.63 | 3 Cenozoic occ. | | PBDB_25406; L_4309; L_5490 | Cope 1867; van Baal et al. 2013; Dutheil 1996 | both | The occ. van Baal et al. is not based on evidence |
| Centroscymnus | 132.6 | 11.63 | | | | | | |
| Chiloscyllium | 121.4 | 15.97 | | | | | | |
| Chlamydoselachus | 86.3 | 0.129 | | | | | | |
| Coupatezia | 83.5 | 33.9 | | | | | | |
| Crassescyliorhinus | 86.3 | 33.9 | | | | | | |
| Cretalamna | 113 | 37.71 | | | | | | Duplicate with *Cretolamna* |
| Cretolamna | 129.4 | 33.9 | | | | | | Records from the Late Eocene based on a poorly preserved tooth (Vasquez and Pimiento 2014) that does not share characters with the genus (Siversson et al 2013), which is unknown above the Early Eocene |
| Ctenopristis | 93.9 | 61.6 | 1 Cenozoic occ. | | PBDB_10050 | Tong and Meylan 2013 | with_evidence | This occ. is not based on evidence |
| Danogaleus | 72.1 | 61.6 | 1 Cretaceous occ. | Reworking (loc. 1155) | PBDB_19793 | Noubhani and Cappetta 1997 | with_evidence | |
| Dasyatis | 132.6 | 0.0117 | | | | | | |
| Deania | 83.6 | 2.58 | 3 Cretaceous occ. | | L_471; L_4734; L_5593 | Muller 1989; Thies and Muller 1993 | with_evidence | Open nomenclature: Reported as *Deania* ? sp. *Deania* ? n. sp. |
| Echinorhinus | 139.8 | 0.0117 | | | | | | |

**Table S4 (continued)**

| | | | | | | | | |
|---|---|---|---|---|---|---|---|---|
| Eorhincodon | 113 | 33.9 | | | | | | Disused and two genus names: *Eorhincodon* Li, 1995 and *Eorhincodon* Nessov, 1999. The latter was replaced by *Pseudomegachasma* Shimada et al., 2015 and the former is considered *nomen dubium* as based on an indetermined carcharhiniform tooth. |
| Eostegostoma | 86.3 | 33.9 | 1 Cretaceous occ. | | PBDB_25978 | Mustafa et al. 2002 | no_evidence | |
| Etmopterus | 72.1 | 0.129 | | | | | | |
| Foumtizia | 72.1 | 33.9 | 2 Cretaceous occ. | Reworking (loc. 1155) | PBDB_19787, PBDB_19788 | Noubhani and Cappetta 1997 | with_evidence | |
| Galeorhinus | 86.3 | 0 | | | | | | |
| Ganntouria | 72.1 | 61.6 | 1 Cenozoic occ. | Reworking (loc. 1155) | PBDB_19783 | Noubhani and Cappetta 1997 | with_evidence | |
| Ganopristis | 86.3 | 61.6 | 2 Cenozoic occ. | Reworking (Dutheil 1996) | L_4300; L_5508 | Dutheil 1996; van Baal et al. 2013 | both | van Baal et al. is considered as "with evidence" whereas no evidence is provided |
| Ginglymostoma | 89.8 | 0.0117 | | | | | | |
| Gladioserratus | 139.8 | 61.6 | | | | | | |
| Gymnura | 100.5 | 2.58 | | | | | | |
| Hemiscyllium | 121.4 | 33.9 | | | | | | |
| Heptranchias | 72.1 | 0.774 | 1 Cretaceous occ. | | PBDB_12248 | Database 2005 | no_evidence | Occurrence assigned to *Xampylodon dentatus* in PBDB |
| Heterodontus | 139.8 | 0 | | | | | | |
| Heterotorpedo | 72.1 | 33.9 | 1 Cretaceous occ. | Reworking (loc. 1155) | PBDB_19799 | Noubhani and Cappetta 1997 | with_evidence | |
| Hexanchus | 132.6 | 0.8 | | | | | | |
| Hispidaspis | 113 | 33.9 | 2 Cenozoic occ. | | L_20698; L_20691 | Li 1997 | with_evidence | |
| Hypolophodon | 72.1 | 5.333 | 1 Cretaceous occ. | | L_29649 | Bogan and Gallina 2011 | with_evidence | |
| Hypolophus | 86.3 | 56 | 2 Cretaceous occ. | | PBDB_25609; PBDB_23909 | Miller 1968; Thurmond and Jones 1981 | no_evidence | Disused genus name, most species previously assigned to this genus are included in the Cenzoic genus *Hypolophodon* Cappetta, 1980 |
| Hypotodus | 83.6 | 7.246 | 2 Cretaceous occ. | | PBDB_12780; PBDB_16378 | Hartstein and Decina 1986; Tulu and Rogers 2004 | no_evidence | |
| Ischyrhiza | 113 | 56 | 3 Cenozoic occ. | Reworked (Smith et al. 1999) | PBDB_23502; L_2108; PBDB_14116 | Cook and Ramsdell 1991; Smith et al. 1999; Van Valen and Sloan 1965 | both | No evidence for Cook and Ramsdell 1991 & Van Valen and Sloan 1965 |

**Table S4 (continued)**

| | | | | | | | | |
|---|---|---|---|---|---|---|---|---|
| Isurus | 93.9 | 0 | 7 Cretaceous occ. | | PBDB_12102; PBDB_12212; PBDB_23926; PBDB_26607; PBDB_1021; L_3675; PBDB_16477 | Chapman 1918; Miller 1966; Armstrong-Ziegler 1978; Bergström et al. 1973; Reeside 1955; Hill 1989; Zangerl and Sloan 1960 | no_evidence | All without evidence except one (Hill 1989) |
| Ixobatis | 72.1 | 61.6 | 3 Cenozoic occ. | Reworking (loc. 1155) | PBDB_13713; PBDB_19778; PBDB_10048 | Hirayama and Tong 2003; Noubhani and Cappetta 1995; Tong and Meylan 2013 | with_evidence | 2 occ. doubled and without evidence |
| Lamna | 132.6 | 0.0117 | 11 Cretaceous occ. | | PBDB_10082; PBDB_12093; PBDB_16049; PBDB_26124; L_29498; PBDB_25292; L_4804; PBDB_26624; PBDB_25656; PBDB_10820; PBDB_10522 | Leriche 1942; Davis 1888; Wieland1896; Giers 1964; Williams 2006; Lucas et al. 1988; Gonzalez-Rodriguez et al. 2013; Bergström et al. 1973; Clark 1893; Thurmond 1971; Malzahn 1979 | both | Old genus name previously used for various lamniform taxa but now restricted to two extant species and their extinct (Cenozoic) relatives |
| Leptocharias | 86.3 | 33.9 | | | | | | |
| Megasqualus | 72.1 | 2.58 | | | | | | |
| Mesiteia | 121.4 | 47.8 | 1 Cenozoic occ. | | PBDB_10599 | Carnevale et al. 2014 | This occurrence is considered dubious as indicated in the reference | |
| Myledaphus | 93.9 | 56 | 2 Cenozoic occ. | | L_3930; PBDB_14163 | Wroblewski 2004; Holtzman 1978 | both | |

**Table S4 (continued)**

| | | | | | | | | |
|---|---|---|---|---|---|---|---|---|
| Myliobatis | 83.6 | 0 | 10 Cretaceous occ. | | PBDB_12328; PBDB_19214; PBDB_23378; PBDB_12776; PBDB_16396; PBDB_24904; PBDB_10068; PBDB_10084; PBDB_12894; PBDB_18083 | O'Leary et al. 2019; Hoganson et al. 2019; Gallagher 1984; Hartstein and Decina 1986; Moody and Suttcliffe 1991; Tabaste 1963; Hartstein et al. 1999; Leriche 1942; Berry 1916; Montgomery and Clark 2016 | | Old genus name previously used for various myliobatiform taxa but now restricted specific extant species and their extinct (Cenozoic) relatives |
| Nebrius | 72.1 | 11.63 | | | | | | |
| Notidanodon | 139.8 | 47.8 | | | | | | The Cenozoic species included in this genus actually belong to the genus *Xampylodon* (see Cappetta et al. 2021) |
| Notidanus | 121.4 | 5.333 | | | | | | Disused (SYN: Hexanchus) |
| Notorynchus | 113 | 0 | | | | | | |
| Odontaspis | 139.8 | 2.58 | | | | | | |
| Onchopristis | 145 | 56 | 1 Cenozoic occ. | Age of occurrence indicated as "Possibly Late Maastrichtian-Paleocene" and "Currently we have poor field records regarding the stratigraphic layer in which this taxon was collected." in O'Leary et al (2019; p. 94) | PBDB_17257 | O'Leary et al. 2019 | with_evidence | |
| Onchosaurus | 93.3 | 47.8 | 1 Cenozoic occ. | | PBDB_26449 | Russell et al. 1988 | no_evidence | Open nomenclature: as cf. *Onchosaurus* sp. |

**Table S4 (continued)**

| | | | | | | | | |
|---|---|---|---|---|---|---|---|---|
| Orectoloboides | 113 | 23.03 | | | | | | |
| Orthacodus | 121.4 | 37.71 | | | | | | Disused and duplicate with senior synonym (*Sphenodus* ) |
| Oxyrhina | 93.3 | 0.0117 | | | | | | Disused |
| Palaeogaleus | 86.3 | 47.8 | | | | | | |
| Palaeohypotodus | 72.1 | 19 | 30 Cretaceous occ. | | | | | Attribution of the Maastrichtian species *Lamna (Odontaspis) bronni* Agassiz, 1843 to *Palaeohypotodus* is not common. Either included *Odontaspis* (Cappetta & Corral 1999) or *Jaekelotodus* (Cappetta 2006) |
| Paraorthacodus | 139.8 | 37.71 | | | | | | |
| Pararhincodon | 113 | 33.9 | | | | | | |
| Parascyllium | 72.1 | 37.71 | 1 Cretaceous occ. | | L_3406 | Siverson 1993b | no_evidence | Unpublished |
| Parasquatina | 93.9 | 61.6 | | | | | | |
| Paratriakis | 93.9 | 61.6 | | | | | | Species with Maa-Dan occurrences (*P. curtirostris* ) belong to another genus (*Palaeotriakis* ) (see Guinot et al. 2013) |
| Plicatoscyllium | 86.3 | 33.9 | 5 Cenozoic occ. | Reworked (loc. 1155 & Dutheil 1996) | L_20696; PBDB_23957; L_5494; PBDB_1005; PBDB_19780 | Li 1997; Thomas et al. 1999; Dutheil 1996; Tong and Meylan 2013; Noubhani and Cappetta 1997 | Both | |
| Pristiophorus | 113 | 0.0117 | | | | | | |
| Pristis | 83.6 | 0.3 | 1 Cretaceous occ. | | PBDB_12929 | Crane (2011) | no_evidence | Open nomenclature: as *Pristis* ? sp. |
| Prosopodon | 72.1 | 61.6 | | | | | | Disused: replaced by *Phosphatodon* |
| Protolamna | 139.8 | 41.2 | 2 Cenozoic occ. | | L_12199; L_28638 | Andrev and Motchurova-Dekova 2010; Reinecke & Engelhard 1997 | with_evidence | |
| Pseudocorax | 113 | 33.9 | 3 Cenozoic occ. | | L_2070; L_2516; L_55 | Li 1997; Yarkov & Popov 1998; Dutheil 1996 | Both | |
| Pseudoginglymostoma | 72.1 | 37.71 | | | | | | |
| Pteroscyllium | 121.4 | 61.6 | 1 Cenozoic occ. | Reworked (loc. 1155) | | Noubhani and Cappetta 1997 | with_evidence | |
| Pucapristis | 72.1 | 63.3 | 1 Cenozoic occ. | | L_14117 | Williamson & Lucas 1993 | with_evidence | Open nomenclature: as *Pucapristis* ? *Standhardtae* + is probably an orectolobiform (see Guinot & Condamine 2013) |

**Table S4 (continued)**

| | | | | | | | | |
|---|---|---|---|---|---|---|---|---|
| Raja | 86.3 | 0 | | | | | | |
| Rhincodon | 93.9 | 0.0117 | 1 Cretaceous occ. | | L_3685 | Hill 1989 | with_evidence | |
| Rhinobatos | 129.4 | 2.58 | | | | | | |
| Rhinobatus | 121.4 | 5.333 | | | | | | Disused and duplicate of *Rhinobatos* |
| Rhinoptera | 100.5 | 0.0117 | 5 Cretaceous occ. | | L_2934; L_29458; PBDB_23377; PBDB_25773; PBDB_19864 | Muñoz-Ramirez et al. 2008; Muñoz-Ramirez et al. 2007; Gallagher 1984; Jain and Sahni 1983; Werner 1989 | both | |
| Rhombodus | 86.3 | 56 | Reworked (Dutheil 1996) | | PBDB_23501; PBDB_13039; L_5512 | Cook and Ramsdell 1991; Solé et al. 2009; Dutheil 1996 | Both | |
| Rhynchobatus | 86.3 | 0.3 | | | | | | |
| Rolfodon | 86.3 | 11.63 | | | | | | |
| Scapanorhynchus | 121.4 | 33.9 | 5 Cenozoic occ. | | PBDB_24487; PBDB_27037; PBDB_24266; L_20348; L_20357 | White 1956; Arambourg and Joleaud 1943; Abbass 1972; Bernardez 1997 | no_evidence | |
| Sclerorhynchus | 93.9 | 56 | 1 Cenozoic occ. | Reworked | L_2102 | Smith 1999 | with_evidence | |
| Scyliorhinus | 139.8 | 3.6 | | | | | | |
| Serratolamna | 86.3 | 33.9 | 10 occ. of *C. aschersoni* included in *Serratolamna* | | | | | Occ. of the Paleogene *Otodus aschersoni* included in *Serratolamna* . The Paleogene species *Otodus aschersoni* is most commonly considered as closer to *Cretolamna* (Cappetta 2006) than to the Cretaceous *Serratolamna* |
| Somniosus | 83.6 | 2.58 | 1 Cretaceous occ. | | PBDB_23174 | Cappetta et al. 2019 | | The Cretaceous occurrence is attributed to *Rhinoscymnus* by the authors, not to *Somniosus* (see Cappetta et al. 2021) |
| Sphenodus | 139.8 | 55.8 | | | | | | |
| Squalicorax | 113 | 56 | 8 Cenozoic occ. | Reworked (Dutheil 1996) | L_3922. L_2274; L_2515; L_550; L_4285; L_430; L_4302; L_4306 | Dutheil 1996; Da Silva 2007; Shourd & Winter 1980; Yarkov & Popov 1998; van Baal et al. 2013 | Both | |
| Squaliodalatias | 100.5 | 15.97 | | | | | | |
| Squalus | 113 | 0 | | | | | | |
| Squatina | 139.8 | 0 | | | | | | |
| Squatirhina | 121.4 | 56 | 2 Cenozoic occ. | Reworked | L_21049; L_21078 | Smith et al. 1999 | with_evidence | |

**Table S4 (continued)**

| | | | | | | | | |
|---|---|---|---|---|---|---|---|---|
| Squatiscyllium | 83.6 | 41.2 | 2 Cretaceous occ. | | PBDB_13869; PBDB_13897 | Fiorillo 1997 | no_evidence | |
| Stegostoma | 72.1 | 33.9 | 2 Cretaceous occ. | | L_30515; L_30526 | Siverson 1993b | no_evidence | Unpublished |
| Striatolamia | 83.6 | 4.9 | 1 Cretaceous occ. | | PBDB_12099 | Chapman 1918 | no_evidence | Cretaceous occurrence represented by *Otodus apiculatus* Agassiz, which is a Cenozoic *nomen dubium* based on a heterogenous series (see Cappetta 2006) |
| Synechodus | 139.8 | 37.71 | | | | | | |
| Triakis | 72.1 | 0 | 1 Cretaceous occ. | Reworked (loc. 1155) | PBDB_19792 | Noubhani and Cappetta 1997 | with_evidence | |
| Youssoubatis | 83.5 | 61.6 | 1 Cenozoic occ. | | PBDB_10051 | Tong and Meylan 2013 | with_evidence | No evidence in that reference |

**Table S5.** K/Pg victim species missing in Data S1 of Gardiner et al. (2026).

| Species name | Author |
|---|---|
| *Angolabatis benguelaensis* | (Antunes & Cappetta, 2002) |
| *Australopristis wiffeni* | Martill & Ibrahim, 2012 |
| *Carcharias heathi* | Case & Cappetta, 1997 |
| *Chiloscyllium broennimanni* | Casier, 1958 |
| *Chlamydoselachus gracilis* | Antunes & Cappetta, 2002 |
| *Cretascymnus quimbalaensis* | Antunes & Cappetta, 2002 |
| *Cretolamna maroccana* | (Arambourg, 1935) |
| *Erguitaia arganiae* | (Arambourg 1952) |
| *Hamrabatis ornata* | Cappetta, 1991 |
| *Hispidaspis turkestanensis* | Zhelezko, 2000 |
| *Ischyrhiza monasterica* | Case & Cappetta, 1997 |
| *Ptychotrygon greybullensis* | Case, 1987 |
| *Scyliorhinus cepaeformis* | Halter, 1990 |
| *Scyliorhinus wardi* | Halter, 1990 |

**Table S6.** Number of species identified as victims (younger age of their last occurrence at 66 Ma), survivors (occurrences distributed across 66 Ma), along with the total number of lineages present in the Maastrichtian (Richness) and corresponding extinction magnitude (%). Data based on Gardiner et al.'s raw dataset and after revision of outlier occurrences of typical Paleogene and typical Maastrichtian species referred to as "Surv. (typical Pg)" and "Surv. (typical K)" here. See Table S2.

| | GEA data | | GEA data revised | |
|---|---|---|---|---|
| | **Total Div.** | **Standing Div.** | **Total Div.** | **Standing Div.** |
| **Richness (D)** | 227 | 109 | 201 | 103 |
| **Victims (*e*)** | 146 | 66 | 188 | 93 |
| **Survivors** | 81 | 43 | 13 | 10 |
| **Surv. (typical Pg)** | - | - | 26 | 26 |
| **Surv. (typical K)** | - | - | 42 | 42 |
| **Extinction %** | 64.32 | 60.55 | 93.53 | 90.29 |

**Table S7.** Number of genera identified as victims (younger age of their last occurrence at 66 Ma), survivors (occurrences distributed across 66 Ma), along with the total number of lineages present in the Maastrichtian (Richness) and corresponding extinction magnitude (%). Data based on Gardiner et al.'s raw dataset and after revision of outlier occurrences of typical Paleogene and typical Maastrichtian genera referred to as "Surv. (typical Pg)" and "Surv. (typical K)" here. Note that 7 genera representing duplicate or invalid names in Gardiner et al. were removed. See Table S4.

| | GEA data | | GEA data revised | |
|---|---|---|---|---|
| | **Total Div.** | **Standing Div.** | **Total Div.** | **Standing Div.** |
| **Richness (D)** | 151 | 120 | 121 | 102 |
| **Victims (e)** | 51 | 37 | 79 | 65 |
| **Surv. (typical Pg)** | - | - | 23 | 23 |
| **Surv. (typical K)** | - | - | 28 | 28 |
| **Survivors** | 100 | 83 | 42 | 37 |
| **Extinction %** | 33.77 | 30.83 | 65.29 | 63.73 |
